# Gaze audits food items for bite points during human withdraw-to-eat movements

**DOI:** 10.1101/2024.07.24.604915

**Authors:** Ian Q Whishaw, Jessica R. Kuntz, Hardeep Ryait, Julia Philip, Jordyn Koples, Jordan Dudley, Jenni M. Karl

## Abstract

Food handling and eating are central to the skill of primate hand movements, and their analysis can provide insights into the evolutionary origins of hand use and its generalization to other behaviors, such as tool use. Vision contributes differently to the reach, grasp, and withdraw-to-eat components of hand use when eating, suggesting that these component movements are controlled by different visuomotor networks with distinct evolutionary histories. This study examines the role of gaze in mediating the withdraw-to-eat movement in human participants eating various food items, including candy, donuts, carrots, bananas, and apples, or pantomiming the eating movements for some of these items. Eye-tracking and frame-by-frame video analyses are used to describe gaze, gaze duration, gaze disengagement, eye blinking, and hand preference in eating each food item. The results show that gaze first identifies points on a food item that the dominant hand can grasp and then identifies points on the food item that the mouth can bite. The hand and finger shaping movements of both the initial grasp and subsequent food handling aid in exposing targets on the food for grasping and biting. The comparison of real and pantomime eating suggests that only real food items possess the affordances that elicit gaze patterns associated with identifying online targets for grasps and bites. The findings are discussed in relation to idea that gaze has a feature-detector-like role linking food cues to the skilled movements of hand shaping to grasp a food item and then to orient a food item to the mouth for biting.

## Introduction

The contribution of vision to hand reaching and grasping in primates, including humans, underlies their ability to exploit a variety of food sources, (Castiello, 1997; Castiello and Dadda, 2019; Churchill et al, 1999; 2006; Karl and Whishaw, 2013; Mitchell et al, 2024; Quinlan and Culham, 2015; Whishaw and Karl, 2019). Investigation of the relationship between vision and hand use in the eating of variegated food thus has the potential of clarifying the origins of the visual control of the hands, including its potential origins in harvesting insects or fruit on the distal branches of trees (Cartmill, 1972, 2012; Sussman & Raven, 1978; Sussman et al., 1978; 2013; Scott, 2019). One approach to understanding the visual control of hand movements suggests that the movements consist of subcomponents, including a reach, a grasp and a withdraw-to-eat, each of which is mediated by a visual neural network (An et al, 2022; Graziano, 2016; Milner and Goodale, 2006; Jennerod, 1999; Karl and Whishaw, 2013; Whishaw et al, 2016). For example, for the reach, visual control is distinctive in that it is proposed to be online and applied to only real target objects and not to reaches for pantomime objects (Goodale et al., 1994; Kuntz et al., 2020). Recent work on the visuomotor adaptions in the feeding behavior of representative members of the three suborders of extant primate suggest further that the withdraw-to-eat movement has subcomponents (de Bruin et al, 2008; Hirsche et al, 2022; Peckre et al, 2023; Whishaw, 2024a,b). A ground-withdraw movement directly transports and releases a food item into the mouth from the location where it was grasped, e.g., from a substrate such as the ground or a tree branch. An inhand-withdraw movement additionally involves holding and manipulating a food item in the hand before bringing the item to the mouth under visual guidance. All members of the three primate suborders direct gaze towards food items to grasp, but only members of the platyrrhine and catarrhine primate suborders additionally use gaze to mediate inhand-withdraw movements. The purpose of the present study was to further examine the properties of the food items that determine whether they are subject to ground-withdraw or inhand-withdraw movements and to examine the contribution of gaze to each type of withdraw movement.

A puzzling aspect of the visual control of withdraw-to-eat movements relates to food size. It might be expected that it could be more difficult both to grasp and then to target a small food item to the mouth. Nevertheless, anthropoid primates, including humans, often disengage their gaze during reaching at about the time, or even before, the grasp of a small food item is completed (de Bruin et al., 2008; Hirsch et al., 2022; Sacrey, 2009; 2011; Whishaw et al., 2024). They then do not use gaze during the withdraw-to-eat movement. This finding suggests that guiding a food item to the mouth can be accomplished using peripheral vision and/or somatosensory cues (de Bruin et al., 2008; Goodman and Tremblay, 2018; Hall et al., 2014; Karl et al., 2012; Edwards et al., 2005). When anthropoid primates hold a large food item during an in-hand withdraw, however, gaze is directed toward the food both during food manipulation and during the initial portion of the withdraw movement. This observation shows that some aspects of food identification and/or manipulation of large food items are dependent on gaze. One reason that gaze might be more closely associated with in-hand withdraw movements could be related to whether a food item protrudes from the hand. The protruding portion might require vision for guidance to the mouth. This hypothesis guided the present study’s examination of the relation of gaze to withdraw-to-eat movement with food items of various sizes, including candy, donuts, carrots, bananas, and apples. In some experiments, participants were asked to pantomime the grasp and withdraw-to-eat movements with a pretend food item, an act that requires both reach and withdraw movements but may not be accompanied by online gaze (Davarpanah et al., 2016; Kuntz et al., 2020; Goodale et al., 1994).

The experiments were performed with adult female and male participants who were video recorded in all experiments and wore eye-tracking glasses during some of the experiments. The frame-by-frame video inspection method of Karl et al. (2018) was used to analyze gaze durations, movement durations, and eye blinks with respect to each reach-to-grasp movement and each withdraw-to-eat movement. The experiments indicated that the use of gaze during food handling and the withdraw-to-eat movement contributes to determining an optimal bite point on a food item,

## General Methods & Materials

### Participants

Participants were 81 right-handed young adults (35 female; mean age 20 ± 0.9 months) recruited from undergraduate and graduate psychology and neuroscience classes at Thompson Rivers University and the University of Lethbridge. Undergraduate students received 1% bonus class credit for their participation. Participant handedness was determined by asking each participant which hand they wrote with. For Experiment 1, 17 right-handed participants (6 female) were drawn from an initial cohort of 23 (12 female), but 6 participants were removed from the analysis due to faulty video recording or eye tracking (Kuntz et al., 2020). Experiment 2 included 16 (female) participants. Experiment 3 included 42 (20 female) participants. Each participant gave informed consent, authorized use of photos or videos for the experimental analyses, self-reported as having no history of neurological, sensory, or motor disorders as well as normal, or corrected-to-normal, visual acuity. The University of Lethbridge and Thompson Rivers University Human Subject Research Ethics Committees approved the studies.

### Experimental Setup

In Experiments 1 and 2 participants wore eye tracking glasses and were tested in a normally lit room with a self-standing height-adjustable pedestal placed in front of them. The pedestal was placed at a horizontal reach distance normalized to the participant’s arm length and the height of the pedestal was adjusted to the participant’s sitting trunk height. When a participant reached with an outstretched arm while seated, they could comfortably grasp an object from the top of the pedestal (Whishaw et al., 2002). In Experiment 3, participants took a seat in a comfortable position on a chair and then selected a food item from a tray, and proceeded to eat the food item.

### Data Collection

#### Video recordings

Video cameras were Sony camcorders with variable shutter speed (HDRCX405) or an iphone 11 (https://www.apple.com/ca/iphone/). Filming was performed at a sampling rate of 30 Hz (1/1000 shutter speed on the camcorders and with the default recording mode on the iphone), with the cameras placed to capture frontal views. In Experiment 2, the frontal eye tracking camera also recorded the participant’s reflection in a mirror, which also provided a participant side view.

Inspection of the videos was performed with Quick time v7.7.7 (https://quicktime.en.softonic.com/mac) or Adobe Premier Pro (2024, https://www.adobe.com/ca) software. The zoom function on Premier Pro was used to confirm eye blinks associated with reaching. The methodology used for video analysis was mainly frame-by-frame video inspection using the method of Karl et al., (2018).

#### Eye movement recordings

Participant gaze and blinks were recorded using a ViewPoint EyeTracker^®^ (Arrington Research, Inc.), a monocular, scene-based, eye-tracking device. Participants wore the eye-tracking glasses for the entirety of the experiment and data was collected at a sampling rate of 30 Hz. A sixteen-point eye calibration was performed prior to data collection and was occasionally adjusted, if necessary, during the experiment if a drift developed between the participant’s gaze-point and the target to be fixated. Visual disengage events were determined by inspecting movements of the participant’s gaze-point within the scene view from the eye-tracking glasses and blinks were determined by inspecting the eye view from the eye-tracking glasses, which recorded the participant’s eye directly.

#### Movement Kinematics

Kinematic analyses of arm, hand, and eye movements were conducted using the open access software program Tracker (https://physlets.org/tracker/). Event times and body and hand coordinates were transferred from Tracker to Microsoft Excel (https://www.microsoft.com/en-ca/microsoft-365/mac/microsoft-365-for-mac) to generate graphical representations and figures in Adobe Illustrator (https://www.adobe.com/ca/products/illustrator.html).

### Behavioral measures

#### General measurement

1. *Total eating time.* Total eating time was the time taken to eat a food item after an experimenter instructed the participant to begin the task to the time that the participant indicated that they were finished eating.

#### Gaze measures

1. *Number of gaze events*. A gaze event was defined as a movement of the head to direct the eyes to a food item or was indicated by the gaze point on the eye tracker (Posner et al., 1987).
2. *Number of ground-withdraw gaze events*. A ground-withdraw gaze event was one in which gaze was directed to the target during the reach -to-grasp.
3. *Number of inhand-withdraw gaze events*. Inhand-withdraw gaze-related events were gaze events directed to a food item that was being held in the hand before being brought to the mouth.
4. *Vicarious gaze events.* Vicarious gaze events were those that were directed to a food item held in the hand that ended with visual disengagement but no withdraw of the hand with the food item to the mouth.
5. *Gaze onset*. Gaze onset was defined as the first video frame on which the participant’s gaze-point fixated on the pedestal/target, indicating that the participant was looking at the target.
6. *Gaze disengage.* Gaze disengage was defined as the first video frame on which the participant’s gaze-point moved away from the target or failed to follow the target as the hand moved to bring a food item to the mouth.
7. *Gaze duration*. Gaze duration was the time between gaze onset and gaze disengage as measured in video frames converted to seconds.

#### Reach measures

1. *Reach*. A reach was a movements of the hand directed to picking up a food item.
2. *Withdraw-to-eat*. Withdraw-to-eat events were defined as individual forearm and/or hand movements that brought a food item to the mouth so that a piece of the food item could taken by the mouth.
3. *Reach duration.* Reach duration was the time from the first video frame that the hand moved to initiate a reach movement to the frame on which the fingers closed to grasp a food item.
4. *Withdraw-to-eat duration.* The duration of the withdraw-to-eat movement was defined as the time from the frame of the first movement of the hand after it grasped the food item to the frame that the food item first touched the mouth.

#### Hand and arm postures

1. *Grasp.* A grasp was a hand movement that purchased a food item. Hand grip postures were scored as precision grips, in which an item is usually held by the phalanx pads of the thumb and one or more of the other fingers, including the phalanx pads or the side of the distal segments of the fingers. Alternatively, they could be scored as power grips, in which an item is held against the hand palm, as defined by Napier (1956). Standard notation of the fingers is used as they are numbered from digit 1 to 5 beginning with the thumb.
2. *Arm posture.* Arm postures when holding a food item were defined with respect to the degree of opening of the elbow. In a hand-up posture, the elbow is flexed, and the forearm and hand are held up and toward the mouth, with the elbow clear of the body or resting on the thigh. In a hand-down posture the elbow and forearm are extended and the hand is resting on the thigh or knee.
3. *Handedness.* When the participant was introduced to the task, they were asked which hand they wrote with, as a measure of self-declared handedness (Corey et al., 2001). When they were holding a food item to bring it to the mouth for eating, the hand used was recorded as either the left or right hand.
4. *Food manipulation events*. A food manipulation event is defined as a change in hand grip on a food item or a change in the hand that held the food. The specific strategy used to orient the food to the mouth, defined by particular changes in grip or handedness, were also scored.

#### Blink counts

1. *Disengage blinks.* Disengage blinks were eye blinks that occurred just as the participant’s gaze shifted from a food item, as indicated by the participant’s gaze-point on the eye-tracker, or via inspection of the videos (Willettet et al., 2023). For experiments in which the eye-tracker was not used, blink occurrence was defined by video inspection, including inspection using the zoon function in Adobe Premiere Pro, which permitted a close-up, frame-by-frame inspection of the eye.

### Statistical Analyses

The data were analyzed using a general linear model repeated-measures analysis of variance (RM-ANOVA) with the statistical program SPSS (v.29.0.1.1). A *p* value of < 0.05 was defined as significant. Group comparisons were made using t-tests or LSD tests. The strength of relationships between independent and dependent variables were indicated by SPSS eta-squared (η^2^) and the strength of main effects was assessed with the SPSS power function. Pearson product-moment correlations were used for data fits and correlations were expressed as r values with associated statistical values. In the experiments, group differences were evaluated for sex differences, which are additionally reported in the experimental results if they occurred.

## Experiment 1: The relation of gaze to reaching for a skittle or donut ball

### Procedure

In Experiment 1, participants reached-to-eat two food items, a skittle or a donut ball (Figure 1A). One group of the participants (n = 7) reached for a round donut ball with a diameter of approximately 29mm and another group of participants reached for skittles, a candy with an approximate diameter of 8mm. Participants were seated in a comfortable upright position with feet flat on the floor and their hands placed in the start position. The start position for the right hand was marked on the dorsum of the right thigh, and participants started with their right thumb and index finger in opposition. The left hand rested in an open and relaxed position on the dorsum of the left upper thigh. Participants then completed a set of practice trials where they reached out and grasped an object and brought it back to their chest. This was done so that participants would be accustomed to the task and to ensure that the recording equipment would not interfere with their reach-to-grasp movements. Participants adopted the start position between trials and waited for a start prompt which was a soft verbal “GO” command from the experimenter. Participants reached-to-eat the food item under two reach conditions, real and pantomime.

1. *Real reach*. In the real reach condition both the pedestal and the food item were present such that the participant reached for a real skittle or donut ball. Participants were instructed to “reach out and grasp the target and bring it back up to your mouth either to eat it or as if you were to eat it”.
2. *Pantomime reach*. In the pantomime reach condition, the pedestal was present but the food item was not. The participant was instructed to “pretend to reach out and grasp the target object as you did when the target was present and bring it to your mouth for eating”.

**Figure 1.**
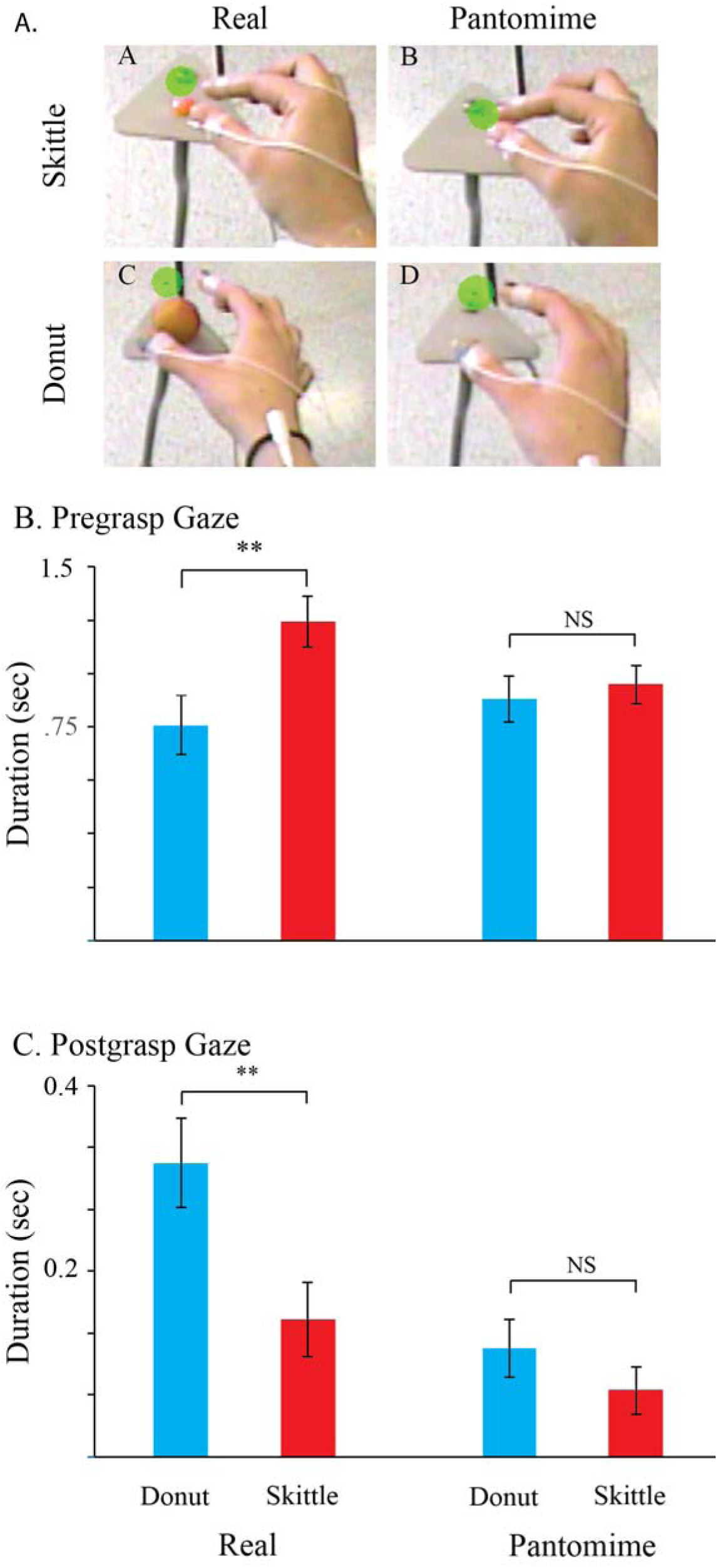
A. The skittle and donut task. Examples of hand shapes at maximum pregrasp aperture (MPA) in the real (left) and pantomime (right) variations of the reaching task. The green dot represents the gaze point provided by the eye tracking glasses. B. Gaze duration associated with reaching for the skittle is longer than that for reaching of reaching for the donut. C. Gaze duration associated with withdraw of the donut is longer than that for withdraw of the skittle. Comparable measures on pantomime reaches gave no similar food-item difference (B,C, Left).

In each context, each participant performed 10 real reaches and 4 pantomime reaches (Karl et al., 2013). The pantomime reaches were always conducted after the participants had completed the real reaches to ensure that all participants were familiar with the real condition before they performed the corresponding pantomime movement.

### Results

When reaching for real targets, the participants’ gaze duration varied according to the movement they were performing and the type of food they were eating. During the reach movement, gaze durations were longer for the small skittle than for the larger donut. In contrast, during the withdraw movement, gaze durations were longer for the donut than for the skittle. There were no comparable differences in gaze durations in the pantomime condition.

*Gaze duration for the reach*. Figure 1B shows that a participant’s gaze duration during the reach was influenced by the type of food item they were reaching for, but only when performing real, but not pantomime, movements. For reach movements, there was a significant interaction between condition and food type, F (1,15) = 8.0, p = 0.013, η^2^ = 0.39, power = 0.49. Follow-up tests revealed that participants displayed longer gaze times when reaching for a skittle than a donut ball in the real condition (p = 0.004), but there was no comparable difference in gaze times relative to food items during the pantomime condition, (p = 0.78). A comparison of reach durations from the beginning of the hand movement to the grasp for the donut and for the skittle showed that reach duration to the donut was significantly shorter than the reach duration for the skittle, t(15) = 3.4, p = 0.004. The main effect of condition was trending towards significant, F(1,15) = 4.3, p = 0.06, η^2^ = 0.22, power = 0.5. The main effect of food type (donut vs skittle gaze time), F(1,15) = 3.96, p = 0.7, η^2^ = 0.21, power = 0.46 was not significant.

*Gaze duration for the withdraw*. Figure 1C shows that a participant’s gaze duration during the withdraw movement was influenced by the type of food item they were reaching for, but only when performing real, not pantomime, movements. For withdraw movements, there was a significant main effect of food type, as gaze duration was longer for the donut ball than for the skittle, F(1,15) = 7.2, p = 0.02, η^2^ = 0.3, p = 0.7. There was also a significant main effect of condition as the gaze durations were significantly longer for the real compared to the pantomime condition, F(1,15) = 16.3, p < 0.001, η^2^ = 0.52, power = 0.96. Follow-up tests showed that the longer gaze duration was only significant for the real condition (p = 0.016) and not the pantomime condition (p = 0.29). The interaction of condition by food item was trending towards significant, F(1,15) = 3.27, p = 0.09, η^2^ = 0.18, power = 0.4. Movement duration for the withdraw-to-eat were not measured because the eye tracking glasses could not follow the hand as it approached the mouth.

## Experiment 2: Small and large carrots

### Procedure

In Experiment 2, participants (n = 16) were fitted with the eye tracking glasses and reached-to-eat a small piece of carrot or a complete carrot with the right hand (Figure 2A). The small piece of carrot was cut from the butt end of a full carrot and had a diameter and length of about 2 cm. The butt end was used so that when oriented to the right side of the pedestal, the grasp size on the carrot piece and the full carrot would be equivalent. The length of the full carrot was about 15 cm, so that when grasped on its butt end, most of the carrot extended away from the hand. The remainder of the experimental procedure was the same as in experiment 1. Participants reached-to-eat the food item under two reach conditions, real and pantomime.

1. *Real reach*. In the real reach condition both the pedestal and the food item were present such that the participant reached for the carrot butt or the full carrot. Participants were instructed to “reach out and grasp the target and bring it back up to your mouth either to eat it or as if you were to eat it”.
2. *Pantomime reach*. A pantomime trial was given after each real reach trial with the pedestal present but the food item removed. The participant was instructed to “pretend to reach out and grasp the target object as you did when the target was present and bring it to your mouth for eating”.

**Figure 2.**
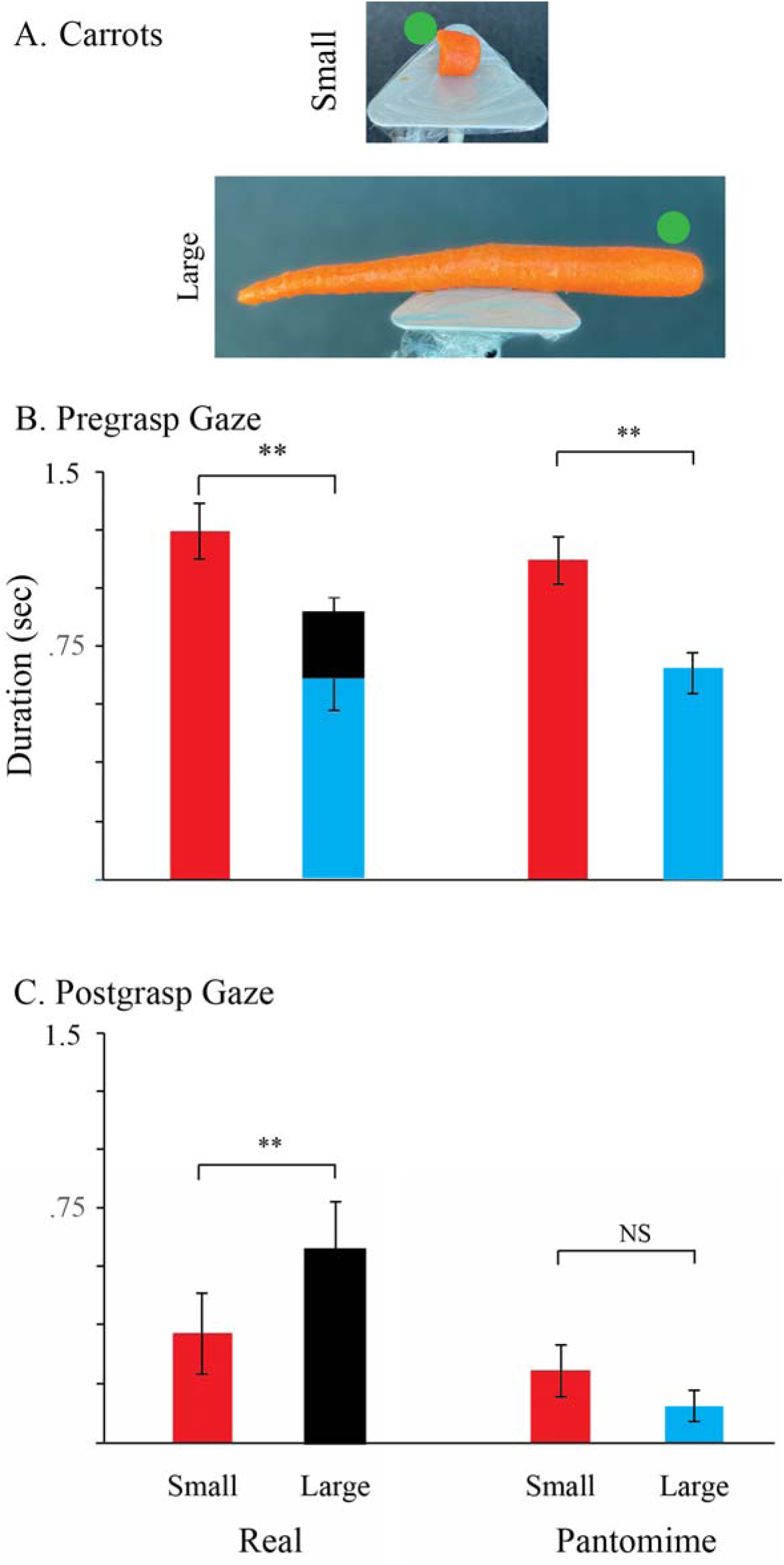
A. The carrot butt and carrot task. Top: the butt end of the carrot is shown along with a calibration ruler. Bottom: the full-sized carrot extends from the pedestal and participants were asked to grasp the butt end. B. Gaze duration prior to and following the real grasps when reaching for the carrot butt and the real carrot. The black portion of the bar for the large carrot is associated with the gaze shift from the but to which the participants were reaching to the far end of the carrot, which occurs before the carrot is grasped. C. Gaze duration following the grasps when reaching for the carrot butt and the full carrot. The black bar represents gaze on the far end of the carrot from the butt end. Note: in the pantomime condition the participants did not shift gaze to the far end of the full carrot.

### Results

During real reach movements participants initially directed their gaze to the butt end of both the small and large carrot. Upon grasping the small carrot, they disengaged their gaze as they began the withdraw-to-eat movement. When reaching for the full carrot, participants shifted their gaze before the grasp was completed from the butt end to the far end of the carrot. Gaze remained on the far end of the carrot as the withdraw-to-the mouth movement began. In the pantomime condition, participants only directed their gaze to the end of the carrot that they grasped.

#### Pregrasp gaze duration

A summary of real gaze durations when reaching to the carrot butt vs a full carrot is shown in Figure 2B. There was a significant effect of carrot size on gaze duration during real reach-to-grasp movements, as the participants visually fixated on the small piece for longer compared to the full-sized carrot, F(1,14) = 7.0, p < 0.019, η^2^ = 0.21, power = 0.08. There was no effect of the test condition (real vs pantomime), F(1,14) = 0.28, p = 0.16, η^2^ = 0.21, power = 0.08 and the interaction between test conditions and food size was not significant, F(1,14) = 2.17, p = 0.16, η^2^ = 0.13, power = 0.30.

The differences in gaze duration related to the size of the target items likely influences reaching time, as the participants reached more quickly for the large carrot in both the real and pantomime conditions. This reach duration difference was likely due to the fact that the full carrot was easier to grasp because its end protruded from the pedestal; i.e., the participants did not have to modify their reach in relation to both the pedestal and the target, Size F(1,14) = 7.02, p = 0.019, η^2^ = 0.33, power = 0.7. There was no significant effect related to the real vs pantomime conditions and no interaction between size and real vs pantomime condition.

#### Postgrasp gaze duration

A summary of real gaze durations when withdrawing-to-eat the carrot butt vs the complete carrot is shown in Figure 2C. There was a significant effect of carrot size on gaze duration during real withdraw-to-eat movements, as participants maintained gaze on the small carrot for a short time after grasping. In contrast, they disengaged from the grasp point on the full carrot before the grasp was complete, F(2,30) = 45.10, p < .001, η^2^ = 0.73, power = 1.0, by shifting their gaze to the other end of the carrot; i.,e., to the end that they would bring to their mouth (the black bar in Figure 2C.

There was a significant main effect of condition, F(1,14) = 18, p < 0.001, η^2^ = 0.56, p = 0.98, because, for the pantomime condition, participants maintained gaze on the butt of both the small and full sized carrot after grasping. The difference in the time of visual disengagement associated, with a longer duration gaze directed to the full sized carrot in the real vs pantomime condition, result a significant condition by food type interaction, F(1,14) = 15.8, p < 0.001, η^2^ = 0.53, power = 0.96.

Participant gaze on the butt end of the full sized carrot was shorter than for grasping the small piece of carrot because they shifted gaze to the far end of the carrot before the grasp. Nevertheless, total gaze time during the withdraw movement (including the visual shift to the terminal end of the full carrot shown by the black bar in Figure 2B) was longer for the full carrot than for the small carrot, t(14) = 9.3, p < 0.001). In addition, the duration of gaze after the grasp was also longer for the complete carrot than the small carrot, t(14) = 2.65, p = 0.019.

#### Blink associated with visual disengage

The eye-tracking glasses collect both the participant’s gaze-point as well as a video of the participant’s eyes, showing that they often blinked when they visually disengaged from the food item as they brought it toward the mouth. Participants blinked during the withdraw-to-eat movement in 81% of real reaches and 79% of pantomime reaches. Figure 3A, shows that there was a significant positive correlation between the time of gaze disengage and the time of a blink during real withdraw-to-eat movements, r(38) = 0.86, p < 0.001. That is, the participants blinked as they visually disengaged from target during real withdraw movement. Figure 3B shows that there was no significant correlation between the time of gaze disengagements and the time of blinks during pantomime withdraw movements, r(37) = 0.22, p > 0.05. In the pantomime condition, participants sometimes blinked as they visually disengaged from the pretend target, other times they simply directed their gaze elsewhere and blinked later.

**Figure 3.**
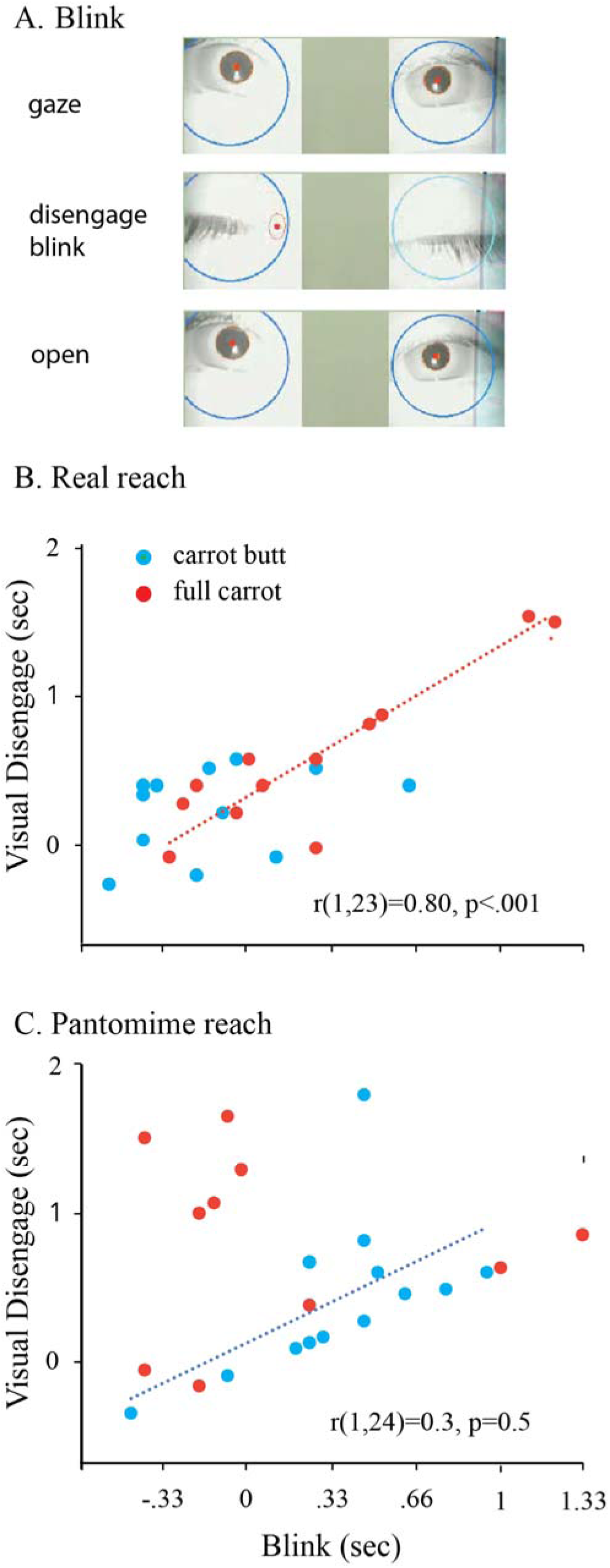
Relation between visual disengage and blinks in real and pantomime reaches for a carrot butt and full-sized carrot. A. Eye view before during and after a blink. (Top, gaze directed toward the full carrot end; Middle, blink; Bottom; gaze redirected away from the carrot.) B. In the real condition, visual disengage was correlated with a blink. C. In the pantomime condition the correlation was smaller and any correlation associated with the small carrot but was associated with gaze directed toward the pedestal.

#### Reach-to-grasp and withdraw-to-eat movement duration

Measures associated with the reach movement are confounded by the placement of the food on the pedestal, as the carrot butt was completely on the pedestal while the full carrot butt extended beyond the pedestal. Measures associated with the withdraw-to-eat movement are also confounded because the full carrot extends beyond the hand, resulting in the hand moving a shorter distance when bringing the full carrot vs the carrot butt to the mouth. The influence of target and placement were was confirmed by a significant condition by movement type interaction, F(1,14) = 4.8, p = 0.047, η^2^ = 0.3, power = 0.5. Follow-up tests indicated that the reach movement was significantly longer in the real compared to pantomime condition (p < 0.02). It was likely easier to grasp the real carrot because it extended away from the pedestal.

## Experiment 3: Carrots, bananas, and apples

### Procedure

In Experiment 3, 10 participants (n = 5 female) ate a carrot (approximately 15cm in length, 1 - 1.5 cm diameter), 11 participants (n = 5 female) ate an apple (7cm diameter), and 11 participants (n = 5 female) ate a banana. Pantomime eating was assessed with 10 participants (n=5 female) Participants brought a friend with them to the test room. The participant was seated comfortably in a chair and took a food item that was offered on a tray. The participant was instructed to eat the food item in a relaxed and normal way, as they ordinarily might, and to freely engage in conversation with their friend or the experimenters while they ate. Participants that selected bananas were instructed to peel the banana as they ate it, rather than peeling it in its entirety first and then eating it. No instructions were given on how to eat the carrot or the apple. Pantomime eating assessment used apples. Participants were told that they would first eat part of and apple and then return it to a plate and then from another plate they would take a mime apple and eat it in the same way that they had eaten the real apple. Eating behaviors were filmed with an iphone mounted on a holder on a table in front of the participant. A waste basket was placed near the participant’s chair, where participants could place the remnants of the food items. Participants were told to begin eating when prompted by the experimenter with the verbal signal, “you can begin”, which was given when they had the food item in hand.

### Results

#### Grip patterns

All participants held the food items using a precision grip in which the thumb tip was opposed to one or more of the fingertips. Participants gripped the food item with the distal phalanx or the sides of the distal portions of the fingers as shown in Figure 4A. Once a participant obtained a preferred grip, there was little change in the grip pattern that they used for the remainder of item eating, although they manipulated the food items by changing finger pressure and orientation as shown in Figure 4B. Only one participant was observed to change grip, and this was to eat the last portions of the apple, during which the participant used both hands to hold the core with pincer grips, between the distal phalanx of the thumb and that of the second finger. All but one participant held food items in the same hand throughout eating. That participant passed a banana from one hand to the other once during eating. Participants also looked at the food items between withdraw movements, during these vicarious gazes they often changed the orientation as illustrated for a participant holding an apple in Figure 4C.

**Figure 4.**
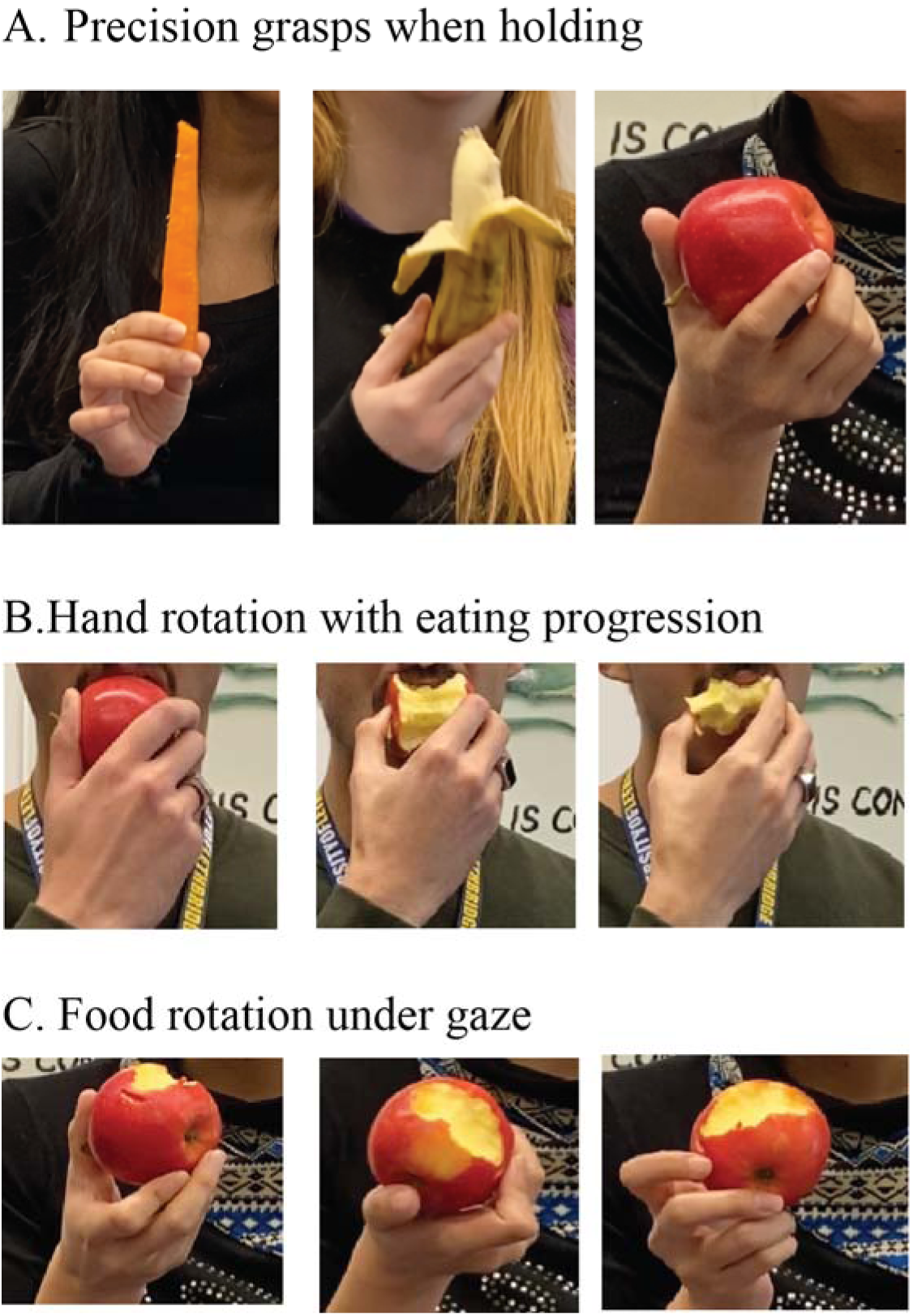
Grasp patterns associated with eating a carrot, banana or apple. A. Precision grasps, involving opposition between the distal thumb pad and the distal pads of the other fingers, were always used for holding the food items. B. Adjustments of the precision grasp pattern as well as wrist rotation are used to position the food objects for mouth placement as eating progresses, as is illustrated for eating an apple. C. With vicarious gaze directed to an apple, a participant rotates the apple to expose target points for biting before the apple is withdraw to the mouth.

#### Handedness

The participants declared that they wrote with their right hand and all but one participant held the food item in their preferred-for-writing hand. One participant held the banana in their left hand and peeled it with their right hand.

#### Arm posture when holding

Figure 5 illustrates postures of the arm used during eating. In one posture, the arm flexed at the elbow with the elbow held up or resting on the lap and the forearm and hand oriented up toward the mouth as shown in Figure 5A. In the other posture the elbow is extended with the forearm resting on the lap and the food item held near the lap. For all participants, the respective postures were used throughout the eating sequences (Figure 5B). More female participants used the elbow flexed posture, whereas more male participants used the elbow extended posture (Chi-square 5.12, p = 0.024).

**Figure 5.**
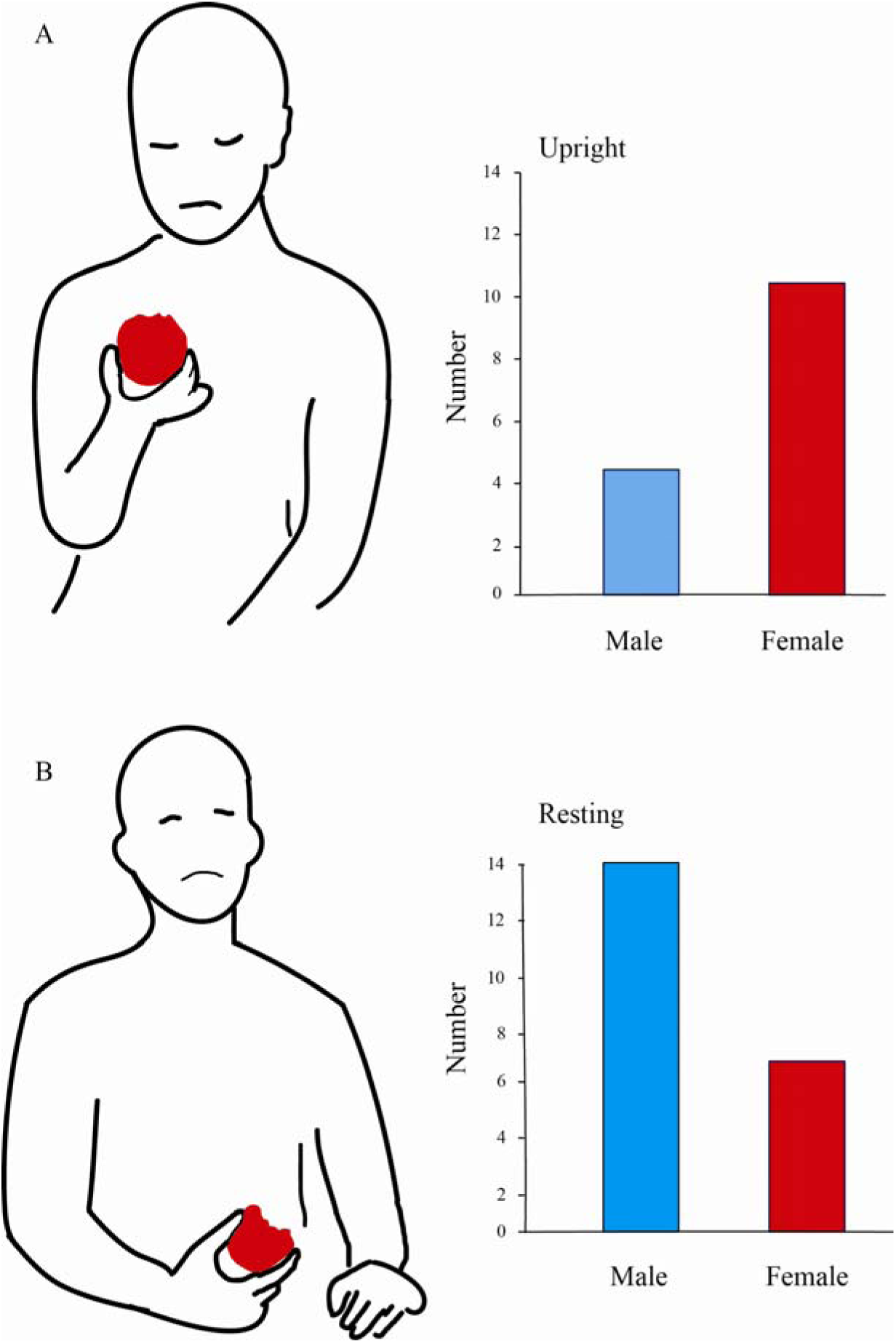
Arm posture for holding a food item. A. The elbow flexed arm posture has the lower arm directed to toward the mouth with the elbow free or resting on a thigh. B. The elbow extended arm posture has the lower arm resting on the lap. Note: more female participants used the flexed arm posture and more male participants used the extended arm posture.

#### Gaze-related withdraw-to-eat

An example of gaze behaviour associated with withdraw-to-eat movements in one participant eating an apple is shown in Figure 6. Figure 6A illustrates the kinematic measures associated with 17 withdraw-to-eat arm movements, with the apple repeatedly brought to the mouth from a holding position with the arm in an elbow-flexed posture. Figure 6B shows the continuous record of gaze movements (red trace) and withdraw-to-eat movements (blue trace), which were closely related and similar for each withdraw-to-eat movement. Figure 6C shows that there were differences in the probability that the participant’s gaze was directed to the food item during a withdraw-to-eat movement depended on the food item that they were eating. When eating carrots, only half of the withdraw-to-eat movements associated were associated with a gaze event. In contrast, when eating a banana or an apple nearly every withdraw-to-eat movement was associated with a gaze event.

**Figure 6.**
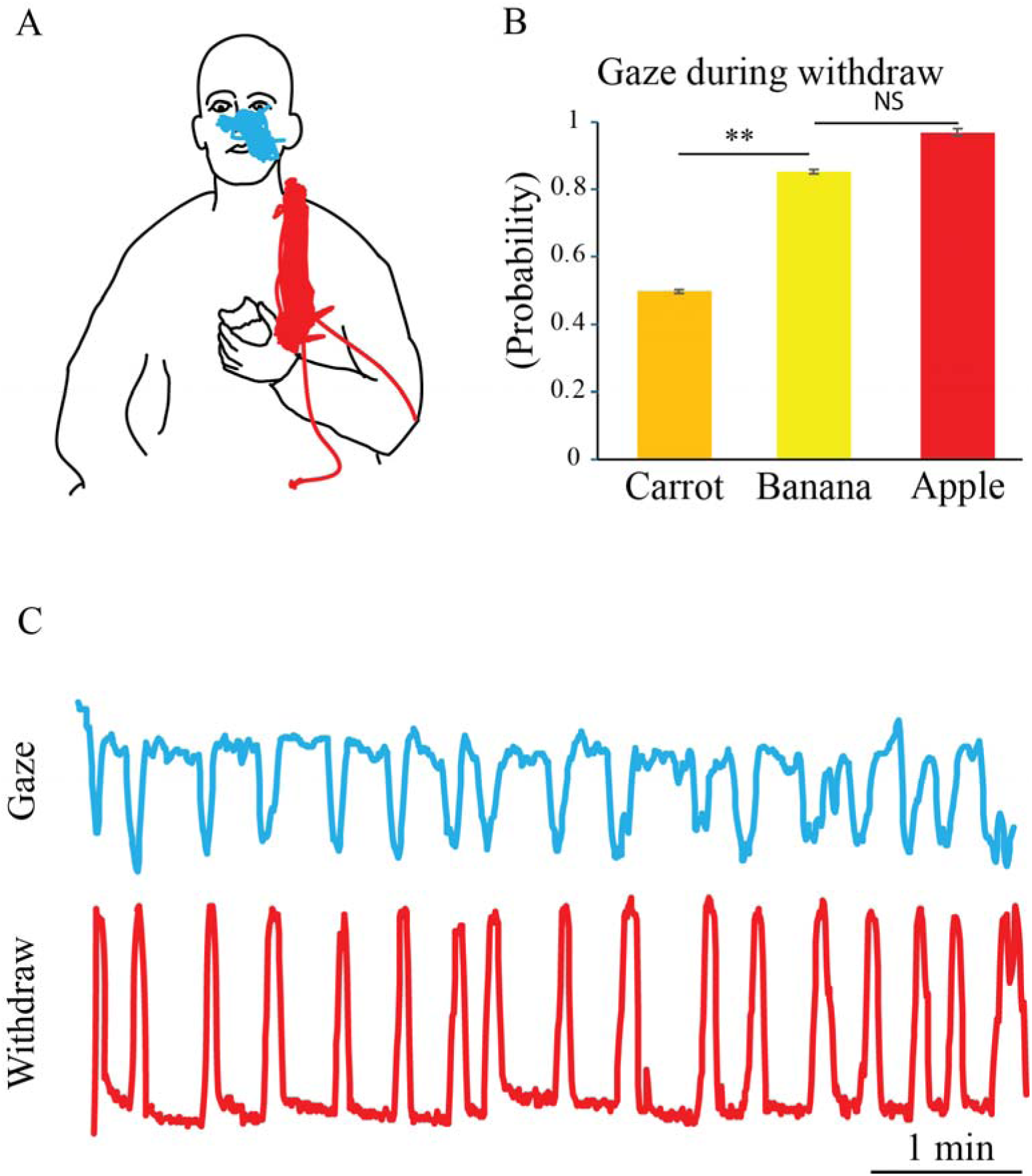
Association between gaze and the withdraw-to-eat movement in a participant who holds an apple in the elbow flexed position. A. Holding posture and kinematic representations of 17 successive head (blue, nose tip tracks) and hand (red, thumb knuckle tracks) of up-down movements taken to eat an apple. B. Relative movement associated with gaze (red, nose tip movement) and withdraw-to eat (blue, thumb knuckle movement) associated with 17 withdraw movements taken to complete apple eating. C. Relationship between the number of gaze events and withdraw-to-eat events in all participants when eating carrots, bananas or apples. Note: gaze is associated with less than half of carrot eating withdraw-to-eat movements but nearly every banana and apple eating withdraw-to-eat movement.

#### Eating duration, gaze duration and gaze type

Figure 7 provides a summary of total eating time, percent of eating time associated with gaze, and percent of gaze events associated with withdraw-to eat movements.

**Figure 7.**
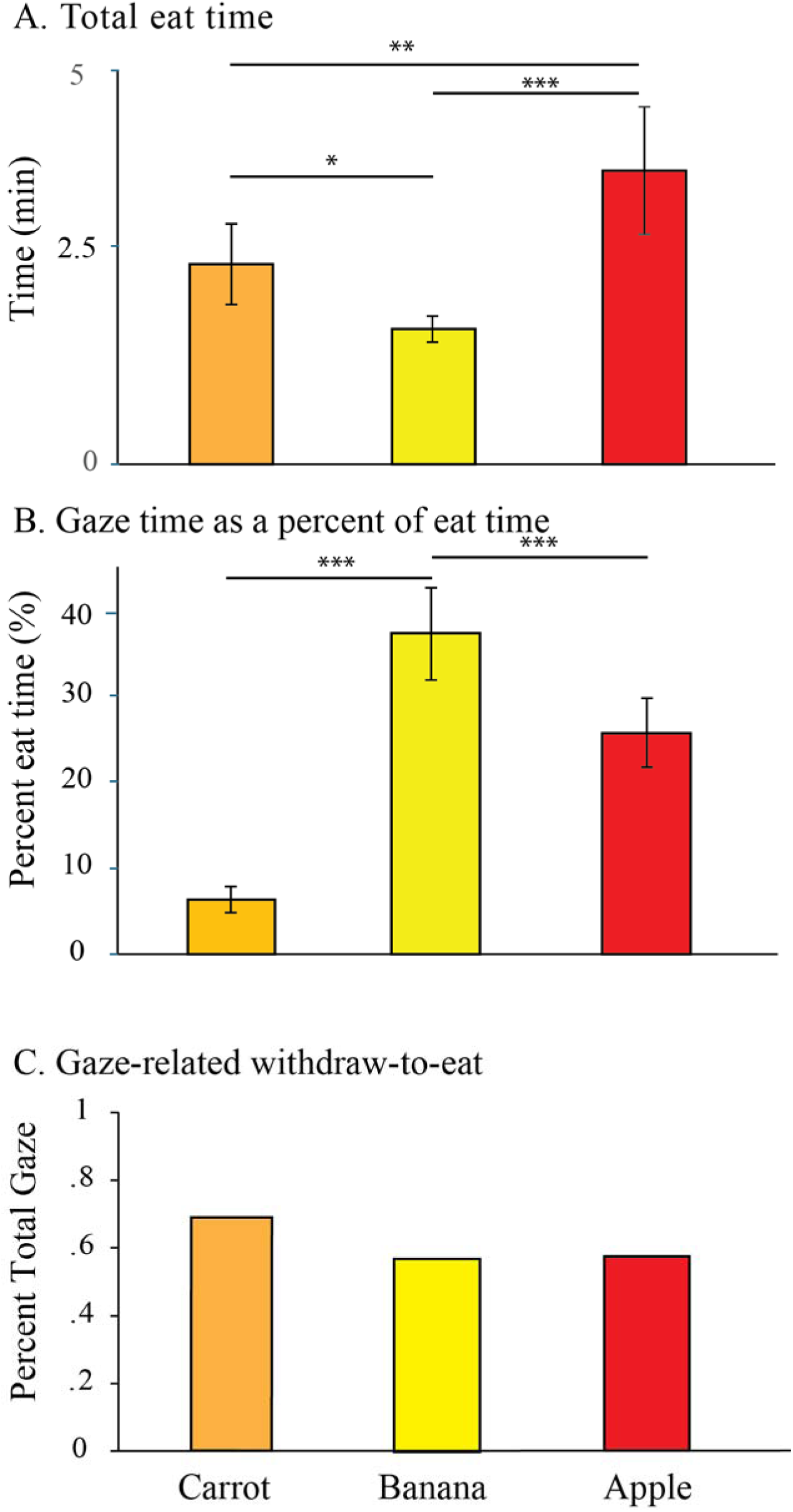
Eating time and gaze times and gaze related withdraw to eat probability. A. Total eating time (mean ± se) to eat carrots, bananas, or apples indicates that it took longer to eat the apple than the other food items. B. Total gaze time as a percent of eating time to eat carrots, bananas, or apples. The long gaze times for the banana were related to peeling. C. About half of all gaze events directed to food items occurred during withdraw-to-eat. The remaining gaze events were vicarious gazes that were not associated with withdraw-to-eat.

*Total eating duration*. A summary of total eating duration for the three different food items is shown in Figure 7A. An ANOVA revealed a significant main effect of food type on total eating duration, F(2,29) = 0.850, p < 0.001, η^2^ = 0.4, power = 0.97. Follow-up pairwise comparisons revealed that participants took longer to eat the apple than the carrot (p = 0.003) and the banana (p < 0.001). Participants also took longer to eat the carrot than the banana (p,< 0.05).

*Total gaze duration*. A summary of total gaze duration as a percentage of total eating duration is shown in Figure 7B. An ANOVA revealed a significant main effect of total gaze duration, F(2,29) = 14.15, p < 0.001, η^2^ = 0.5, power = 1. Follow-up pairwise comparisons showed that participants spent a significantly greater proportion of time visually fixating on the banana relative to the carrot (p < 0.001) and apple (p < 0.003). This is most likely related to the time spent peeling the banana, an activity that was performed almost entirely under gaze. Participants also spent a significantly greater proportion of eating time visually fixating on the apple relative to the carrot (p < 0.001), likely reflecting the fact that some participants seldom looked at the carrot during eating.

*Gaze related withdraws*. Gazes directed toward a food item could be divided into two types. Vicarious gazes were gazes directed toward a food item that was then not withdrawn to the mouth. Withdraw-to-eat gazes were directed toward a food item that was then brought to the mouth for eating. Figure 7C shows the proportion of withdraw-to-eat gazes as a percent of all gazes. About half of all gazes were withdraw-to-eat gazes and an ANOVA did not give a significant main effect of food type associated with the percent of gaze associated withdraw-to-eat events, F(2,29) = 2.1, p > 0.05, η^2^ = 0.

#### Gaze withdraw duration related to food inspection

Figure 8 depicts the correlation between gaze duration and total withdraw-to-eat time for each of the food items. Gaze duration included time spent looking at the food item as it is held plus time spent looking at it as it is withdrawn to the mouth (the first portion of the withdraw movement). Total time of the withdraw-to-eat included gaze time plus the period of withdraw after visual disengage during which a food item brought to be placed in the mouth and positioned for biting. T\correlations in Figure 8 show that the variation in gaze time is due mainly to the time spent looking at the food item before the withdraw movement begins. As illustrated in the Figure 8A, the correlation for the carrot is relatively low, as gaze was not associated with many withdraw-to-eat acts. The correlation for the banana (Figure 8B) and the apple (Figure 8C) are high as gaze was associated with most withdraw-to-eat acts. The insert in Figure 8 (top) depicts the mean ± se of the gaze time vs. total withdraw durations of each of the participant’s correlation for each food item. A comparison of correlation for individuals as a function of food item was significant, F(2,29) = 9.85, p < 0.00. Pairwise comparisons confirmed that the strength of this relationship was weaker for the carrot, compared to the banana (p = 0.016) and apple (p = 0.01), which were not different.

**Figure 8.**
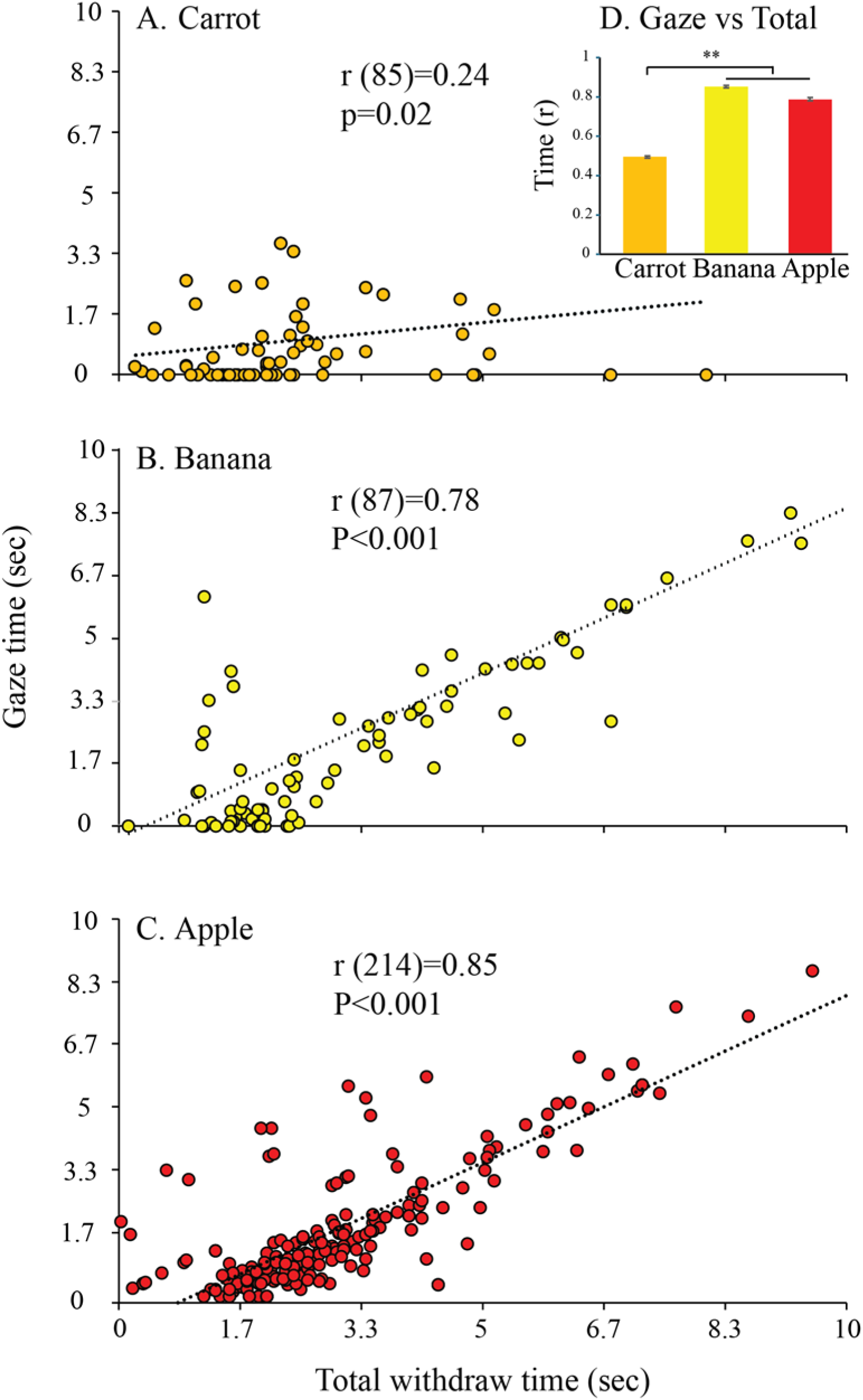
Correlations between gaze duration and total withdraw-to-eat time. A-C. Group results for participants eating carrots, bananas, or apples. D. Insert bar graph shows the correlations (mean±se) averages obtained from individual participants eating each food item. Note: Positive correlations show that the duration of gaze on the food accounts for most of the variation in the total duration of withdraw-to-eat time. The nonsignificant result for the carrot results from many carrot withdraw-to-eat movements being made without associated gaze, but there no withdraw-to-eat events associated with apple eating that were not associated with gaze, however brief.

#### Blink events related to gaze disengage

When a participant who had directed gaze to a food item then brought the food item toward the mouth, the visual disengage was often associated with a blink. Figure 9 (top) illustrates this relationship. The participant first looks at a food item as it is held on the lap and then maintains gaze on the food item during the withdraw-to-eat until gaze disengage, which occurs before the food reaches the mouth (Figure 9 A-C). Figure 9D shows kinematic traces illustrating the timing and the duration of a blink in relation to the withdraw-to-eat movement. The blink occurs just before the hand reaches the mouth, at about the same time that gaze disengage occurs and about the same time that the head is raised so that the food can be accepted into the mouth.

**Figure 9.**
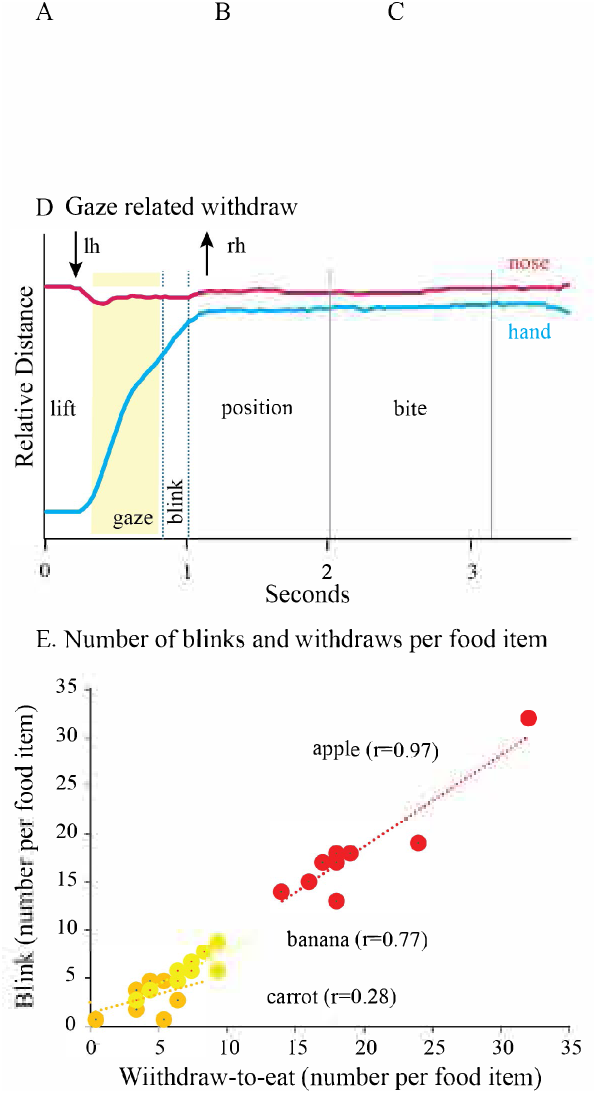
Relation between visual disengage and a blink during apple eating. A-C. For the withdraw; the apple rests in the hand in the elbow extended position, is withdrawn toward the mouth, and placed in the mouth. D. Kinematic trace of the nose tip (red) indicate head movement, to look toward or away from the food item, and the corresponding trace of the knuckle of the thumb (blue) represent hand movement to bring the apple to the mouth. Note: the withdraw is divided into three phases: the lift to bring the apple to the mouth, the positioning of the apple in the mouth, and then a bite in which a piece of apple is taken. Gaze is associated with the lift. Disengage is associated with the blink and occurs before the apple reaches the mouth. E. Correlations between withdraw movement associated with gaze and the occurrence of a blink for participants eating a carrot, banana or apple. (lh - head lower, rh – head raise).

Figure 9E shows the relationship between the number of blinks and the number of withdraw-to-eat movements associated with gaze. The correlation for the carrot was not significant, F(1,8) = 2.3, p = 0.16, as many carrot withdraw-to-eat movements were not associated with gaze directed to the carrot. Correlation were significant for the banana, F(1,10) = 17.8, p<0.002, and the apple, F(1,8) = 48.4, p<0.001.

When participants made vicarious gazes to a food item, gaze disengagements from that item were also associated with a blink. A correlation that included all vicarious gaze events to food items that did not involve a withdraw to the mouth, showed that the correlation (r=0.91) between disengage and the occurrence of a blink, was significant F(1,30) = 36.7, p<0.001.

#### Pantomime apple eating

After a participant had made a first withdraw-to-eat with a real apple or the pantomime apple, the next three withdraw-to-eat movements were examined for the presence of gaze associate with the withdraw-to-eat. For the real apple, gaze was directed to the apple on 29/30 withdraw-to-eat occasions *vs* 13/30 for the pantomime condition, a difference that was significant as shown by a t-test for paired samples, t(9) = 3.1, p = 0.006.

### Discussion

The present study investigated the role of gaze during hand withdraw-to-eat movements of human participants bringing food items of varying sizes and types to the mouth. The hypothesis was that if food items are large and protrude from the hand, gaze would be employed to localize and guide the protruding portion of the food item to the mouth. The experiments show that gaze is not engaged by size *per se* but by the determination of how a food item is to be received by the mouth and bitten. The results suggest that gaze serves a feature-detector-like function in determining the affordance offered by food, first to influence hand movements for grasping the food and then to determine mouth movements for grasping the food as it is brought to the mouth. The role of gaze in determining mouth grasp points on food items and the assistance provided by the hands in exposing grasp points, suggests that withdraw-to-eat movement is a candidate behavior that has contributed to the evolution of the visual control of hand skills in primates more generally.

Human participants employ two types of withdraw-to-eat movements to bring food grasped in the hand to the mouth. A ground-withdraw movement directly transports and releases a food item into the mouth from the location where it is grasped, e.g., from a substrate, such as a pedestal in the present study. An in-hand-withdraw movement involves holding and manipulating a food item in the hand before bringing it to the mouth. The results of the first two experiments support the idea that food size, and the extent to which a food item protrudes from the hand, engages gaze. In the first experiment, the comparison of smaller and larger food items in real and pantomime reaching conditions showed that gaze duration is scaled to the size of food items with smaller items invoking longer gaze during the reach and larger items invoking longer gaze during the withdraw. Similar scaling of movement speed and gaze duration did not occur in pantomime conditions (see also, Coats et al., 2008; Goodale et al., 1994; Jeannerod, 1991; Kennedy et al., 2015; Kuntz et al., 2020; Plamondon and Alimi, 1997; Quinlan and Culham, 2015). The second experiment investigated whether the extent to which a food item protruded from the hand influences the use of gaze during the withdraw-to-eat. Participants reached for a carrot butt or a whole carrot. When grasped on its end, the whole carrot protruded appreciably from the hand. When reaching for the whole carrot, all participants shifted gaze from the grasp point, before the grasp was completed, and directed gaze to the far end of the carrot, the end that was to be placed in the mouth. A similar gaze shift was not associated with a pantomime reach for a whole carrot. The gaze shift directed to the portion of the carrot that protruded from the hand was also associated with a long gaze duration than for the carrot butt, suggested that gaze is used to identify the protruding portion for subsequent eating.

The third experiment, in which participants spontaneously ate a carrot, banana, or apple all of which protruded from the hand, suggests that it is not the size or protrusion of an item from the hand but rather the bite points on a food item that determine the use of gaze. Participants were much less likely to use gaze to bring a carrot or banana to the mouth even though they extended appreciably from the hand, but always used gaze when withdrawing an apple, which if anything, protruded somewhat less. This may because the carrot and banana present no ambiguity with respect to how they are to be presented to the mouth, but for apple eating, each bite on the apple was directed to the margin of the previous bite. Thus, the progression of apple eating required continuous use of the hands for reorienting the apple under gaze to present the next bite point to the mouth. It was clear from the way that the participants ate the carrot and the banana that determining a bite point on the ends of the items was not necessary, as a bite was always directed to the item’s end. We found support for the relevance of bite points to the use of gaze in our video archive, which included four people eating popsicles, a food item that is held by a stick and which also protrudes from the hand. All participants gazed at the popsicle when picking it up, but three of the participants never looked directly at the popsicle again despite bringing it to the mouth many times. Possibly, for food items that lack ambiguity with respect to how they are taken by the mouth, the location and guidance of the end are determined using peripheral vision and/or somatosensory contact with the mouth.

Participants also made vicarious gazes to the food items as they held them, usually during interbit intervals when they were chewing, and often manipulating the item as they did so. Again, there were more vicarious gazes directed to the apple than to the carrot and banana, suggesting that vicarious gazes are involved in planning subsequent withdraw-to-eat movements. These gazes may comprise a vicarious nonconscious procedural audit of the food item. It is interesting that Redish (2016) has suggested that vicarious actions more generally are associated with assessing the possibilities in forthcoming action. The participants were not asked to pantomime eating an apple, the movements these eating movements were associated with reduced gaze frequency. Movement and duration scaling associated with real vs pantomime reaching has been taken to indicate that the real movements are online action (Goodale et al., 1994) and the similar finding obtained here suggest that the visual duration scaling to withdraw-to-eat movement suggest that it is similarly an online action.

Previous studies have noted that humans and some other representative anthropoid primates blink at about the time they visually disengage when grasping a food item or during the withdraw-to-eat movement (de Bruin et al., 2008; Hirsch et al., 2022; Whishaw et al., 2024). These blinks have several suggested causes including; relief from the accommodation required to visualize the target, protecting the eyes from the item when bringing the hand to the mouth, switching from visual to somatosensory guidance, or reflecting a central network change such as alternating between an attention network and the default network (Ang and Maus, 2020; Brych and Handle, 2020; Jaschinski et al., 1996; Nakano et al., 2013; Willett et al., 2023). Here we made a new observation of the relation between blinks and gaze that has a bearing on these suggestions. Blinks occurred when the participant visually disengaged from vicarious gazes, gazes that were not associated with a withdraw-to-eat. This observation weakens the idea that the function of blinks is to protect the eyes from potential injury that could be inflicted by food items brought toward the face. It seems likely, therefore, that blinks are associated with an attentional shift or the removal of visual attention from a target (Daza et al., 2020; Sakai et al, 2017). Interestingly, Irwin (2011) suggests that an eye blink follows an attentional shift, which might suggest that visual attention directed to the food item inhibits blinking and that an attentional shift associated with gaze disengage provides an opportunity of a blink to occur.

There were other notable features of hand use related to the eating of various types of food. First, participants reached-to-grasp and withdrew-to-eat predominantly using the hand with which they wrote. Only one participant was observed to eat a banana with their declared nonpreferred hand. This observation is consistent with the strong relation between handedness for eating and handedness more generally (Sacrey et al., 2013). The observation is relevant to comparing the handedness in eating by nonhuman primates to that of humans, in which only a small number of subjects are usually available for a study (Fagot and Vauclair, 1991; MacNeilage et al., 1987; Salmi et al., 2023). Second, the participants used precision grasps for all grasping, holding, and eating acts. They not only grasped and held small food items with various precision grips between the thumb and the proximal fingers (Wong and Whishaw, 2004), but they also mainly held the apples, which in terms of diameter were the largest food items, in a similar way. Most participants held the apple between the thumb and middle finger and rotated the apple to orient it for eating using a combination of the index finger manipulation and rotatory movements of the hand. Thus, an entire apple was consumed with little to no change in the basic grip but with many changes in its pattern. This observation suggests that grip types in holding (Elliott and Connolly, 1984; Feix et al., 2016; Macfarlane and Graziano, 2009; Napier, 1956; Pouydebat et al., 2009; Spinozzi et al., 2004; Whishaw et al., 2024) likely serve the function of cooperating with vision in the determination of bite points. The observation of the participants use of precision grips is consistent with Napier’s (1956) suggestion that it is function and not size of an object that determines a grip choice. For inhand-withdraw food holding, function would be to expose a bite point, and possibly other mouth-related handling points. Finally, we observed only one sex difference in our analysis of eating-related gaze behavior, which was the more frequent posture of food holding with a flexed, as opposed to extended, elbow by female participants. This may be an example of one of many socially-related sex differences in behavior (Becker et al., 2005; Wood and Eagly, 2007).

In conclusion, the results of the present study suggest that gaze has a feature-detector-like role (Lindeberg, 1998) in determining bite points on food items as they are brought to the mouth for eating. This observation expands the recognized role of gaze from determining a hand grasp point on a food item to also determining a mouth grasp point on a food item. Previous studies have noted that objects can be grasped in relation to their end point comfort or end point use (Martin, 1994; Rosenbaum, 2012), but the present study shows that for food eating, further gaze analysis is necessary for the interactions that the mouth may have with the food item. It is noteworthy that for some food items, gaze was not associated with withdraw-to-eat, suggesting that some aspects of food handling by the mouth may rely on peripheral vision or the interaction of somatosensory cues related to the hands and mouth. Finally, that there interdependences between gaze, hand movement, and mouth movement is consistent with the suggestion that they may be mediated by an oromanual motor cortex area dedicated to food eating (An et al., 2022; Graziano, 2016; Karl and Whishaw, 2013).

## Acknowledgements

This work was supported by the Natural Sciences and Engineering Research Council of Canada (NSERC) [JRK], NSERC Discovery Grant [JBD & JMK]. The authors would like to thank Tsz Yin (Ian) Fung for his help with data collection, Kalob Barr for his help with data analysis, and Ali Mashhoori for his expertise and assistance with MATLAB.

## Conflict of interest

The authors declare no competing financial interests.

## Notes

### Competing Interest Statement

The authors have declared no competing interest.

